# Metabolic syndrome enhances SARS-CoV-2 disease severity and reduces mRNA vaccine efficacy in a mouse model

**DOI:** 10.1101/2025.01.15.633120

**Authors:** Elizabeth Geerling, E. Taylor Stone, Danielle Carpenter, Alexandria M. Dickson, James D. Brien, Amelia K. Pinto

**Affiliations:** Foundation for the National Institutes of Health, North Bethesda, MD; HDT Bio Seattle, WA; Department of Pathology, Saint Louis University School of Medicine, Saint Louis, MO; Department of Molecular Microbiology and Immunology, Saint Louis University School of Medicine, Saint Louis, MO; Department of Microbiology, Immunology, and Molecular Genetics, University of Kentucky, Lexington, KY 40508

**Keywords:** MetS, metabolic syndrome, obesity, SARS-CoV-2, viral infection, vaccine efficacy

## Abstract

Metabolic syndrome (MetS) is a cluster of pathophysiological conditions linked to the disruption of metabolic processes associated with energy storage and consumption. Approximately one-third of adults in the United States are currently diagnosed with MetS. Patients with MetS experience higher mortality rates following SARS-CoV-2 infection and exhibit poor vaccine efficacy following influenza virus vaccination compared to metabolically healthy individuals. However, the specific impact of MetS on immune responses to SARS-CoV-2 infection and vaccination has not been widely studied. To address this gap, we utilized high-fat diet feeding to establish a murine model of MetS, in which mice exhibit the same diagnostic criteria for MetS as human patients. We then used high-fat diet-induced MetS mice and regular chow diet-fed wild-type mice to analyze immune responses to SARS-CoV-2 infection and vaccination. Following SARS-CoV-2 infection, we monitored mice for disease severity, measured levels of virally induced inflammation, and quantified viral titers in various tissues. Our results indicate that MetS mice exhibit accelerated mortality post-infection, accompanied by elevated mRNA levels of inflammatory cytokine transcripts at sites of infection. Additionally, MetS alters the degree of viral replication across various tissues. Furthermore, our vaccination studies revealed that MetS reduces the potency of vaccine-induced neutralizing antibodies against both the ancestral SARS-CoV-2 strain and the Delta variant. Overall, our findings suggest that MetS exacerbates SARS-CoV-2 disease severity and diminishes vaccine efficacy, underscoring the need for tailored strategies to protect individuals with MetS from severe outcomes following infection and vaccination.

**Impact:** This study represents the first detailed account of the failure to generate protective neutralizing antibody responses to SARS-CoV-2 in mice with metabolic syndrome (MetS) following vaccination. The insights gained from this research inform future vaccine design and aid in identifying individuals at greater risk of breakthrough infections. By specifically isolating the risk associated with MetS, rather than obesity alone, these findings lay the groundwork for future investigations aimed at enhancing immune responses in individuals with MetS. This work is highly relevant to a broad audience, as it addresses a critical unmet need in vaccine development and provides essential guidance for the rational design of vaccines to protect vulnerable populations affected by MetS.

## Introduction

Emerging in late 2019, the novel coronavirus, severe acute respiratory syndrome coronavirus-2 (SARS-CoV-2), has caused at least 619 million cases worldwide, and is responsible for at least 6.55 million global deaths [1]. Given its high transmission rate and propensity for inducing coronavirus disease 2019 (COVID-19) in infected individuals, SARS-CoV-2 has put a dramatic strain on public health [2, 3], especially when coupled with the rising incidences of Metabolic syndrome (MetS) diagnoses. Human cohort studies have revealed that MetS is correlated with heightened SARS-CoV-2 viral disease severity [4-7], with studies indicating that MetS is a better prognostic indicator for severe SARS-CoV-2-induced disease than its individual components, like obesity [8]. Further compounding the complexity and toll of the SARS-CoV-2 pandemic, this virus has mutated over time, resulting in the emergence of numerous variants of concern. A viral variant of concern harbors at least one amino acid mutation which confers a greater degree of human transmissibility, resistance to anti-viral treatments, or diminished vaccine efficacy [9]. Around October of 2020, the SARS-CoV-2 B.1.617.2 (delta) variant was first identified, and quickly became the dominant circulating strain. This variant became quickly widespread due to its enhanced transmissibility, shorter incubation period, and higher basic reproductive number when compared to the ancestral strain [10-13]. Alarmingly, chances for severe COVID-19 following infection with the B.1.617.2 variant were higher compared to the ancestral SARS-CoV-2 strain spiked, with one retrospective cohort study showing that when compared to ancestral SARS-CoV-2 infections, B.1.617.2 variant-induced infections increased hospitalization rates by 108%, intensive care unit (ICU) admissions by 235%, and death rates by 133% [14]. As the B.1.617.2 variant accumulated mutations that allowed it to escape some degree of vaccine-conferred immunity, breakthrough infections among fully vaccinated individuals became common amidst the emergence of this viral variant. Interestingly, a large portion of individuals who experienced such breakthrough infections harbored at least one MetS-associated comorbidity [15-17].

MetS is a pathophysiological disorder that is diagnosed as a result of aberrant metabolism in an individual. MetS etiology varies and is multifactorial, including characteristics like weight gain, insulin resistance, sedentary lifestyle, genetic components, and advanced age [18]. To be diagnosed with MetS, an individual must concurrently experience at least three of the following disorders: hyperglycemia, dyslipidemia, hypertriglyceridemia, abdominal obesity, or hypertension [18-20]. Unfortunately, MetS cases are projected to continually rise, and have already risen over 35% since 1988, resulting in one-third of United States adults currently diagnosed with this syndrome [18]. In addition to the criteria used to diagnose MetS cases, these patients often concurrently experience non-alcoholic fatty liver disease (NAFLD), insulin resistance, and chronic inflammation. Persistent inflammation is believed to be the underlying link between these MetS-associated conditions [21]. Prior studies have tried to link obesity with increase disease susceptibility and poor vaccine responses. However obesity alone is a poor predictor of vaccine responses [22] as metabolically healthy obese individuals are not more susceptible to disease [23] and individuals with MetS who are not obese have increased susceptibility to disease. Independently, each MetS-associated comorbidity has been linked to enhanced viral disease severity based on retrospective human cohort studies [24-30]; these findings are alarming to note as they highlight that when these conditions present together, as occurs in the MetS state, the likelihood for MetS patients to experience severe viral disease outcomes increases. In addition to blunted immune responses following viral infection, some studies have addressed whether increasing vaccination rates of individuals with MetS could provide protection against enhanced viral disease severity. These studies revealed that MetS also blunts hepatitis B, influenza virus, tetanus, and rabies vaccine-conferred immunity (reviewed in [31]). The mechanisms driving MetS-induced viral and vaccine-conferred immune defects are unknown, and further investigation is warranted to determine if therapeutic targets can be identified to better protect individuals with MetS from aberrant antigen-specific immunity.

In the current study, we utilized a murine model of MetS to interrogate the impact of metabolic dysfunction on immunity to SARS-CoV-2 infection and vaccination. Following SARS-CoV-2 infection, our results indicated that MetS mice experience accelerated mortality when compared to wild type counterparts, in addition to higher inflammation sites of infection. MetS also impacted viral burden at infection sites. Further, our vaccination studies led us to conclude that MetS impaired the neutralization capacity of vaccine-conferred antibodies against both the ancestral SARS-CoV-2 strain, as well as against the B.1.617.2 variant. Taken together, our results support the findings that MetS enhances SARS-CoV-2 viral disease severity and reduces vaccine efficacy. These findings highlight a previously underappreciated link between MetS and impaired adaptive immune responses, contributing to reduced vaccine efficacy. From a broader global health perspective this study emphasizes the need to consider MetS when designing vaccine strategies.

## Results

### High fat diet feeding leads to pathophysiology indicative of metabolic syndrome

To interrogate the impact of MetS on immune responses to SARS-CoV-2 infection and vaccination, we developed a murine model of MetS based upon our previous studies utilizing a diet-induced obesity mouse model of West Nile virus (WNV) infection [32]. 3-week-old female and male C57BL/6J (B6) or B6.Cg-Tg(K18-ACE2)2Prlmn/J (K18) mice were fed either a regular chow diet (wild type) or a high fat diet. In our previous work, we noted that mice fed the high fat diet weighed 25% more than wild type (WT) counterparts after approximately 12 weeks [32], at which point these mice were considered to be obese as supported by previous literature [33, 34]. In our current study, we expanded this obesity model to examine the mice for indicators of MetS as more recent findings have revealed the importance of considering overall metabolic health rather than obesity alone [35]. Further, MetS has been correlated with heightened viral disease severity in SARS-CoV-2 patients [4-7], with one study indicating that MetS is a better prognostic indicator for severe SARS-CoV-2-induced disease than its individual components, like obesity [8]. To test mice for signs of MetS, serum was collected from WT and high fat diet-fed mice at 12 weeks post-diet administration. Serum levels of cholesterol, glucose, and triglycerides were measured to determine if these mice displayed elevated levels of these MetS diagnostic criteria (**Figure 1**). In **Figure 1A-C**, we noted that serum levels for each of the following MetS diagnostic criteria were significantly elevated in high fat diet-fed mice when compared to WT counterparts: cholesterol (p=0.0003), glucose (p=0.0003), and triglycerides (p=0.0011). These findings confirmed that the high fat diet-fed mice in our murine model exhibited elevated levels of each criteria used to diagnose MetS that we tested: obesity [32], hyperglycemia, hypertriglyceridemia, and hypercholesterolemia.

**Figure 1:**
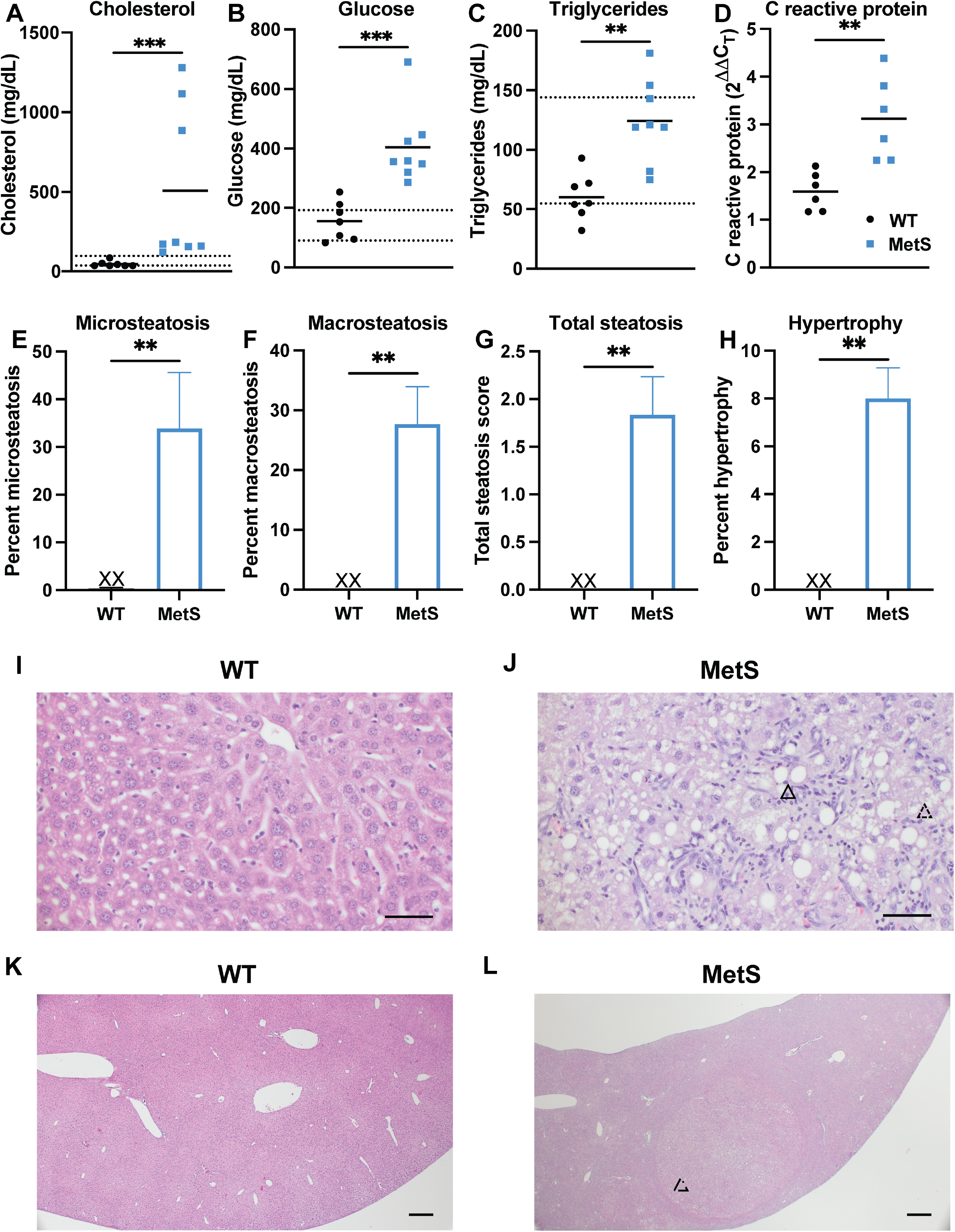
High fat diet feeding leads to pathophysiology indicative of metabolic syndrome. (A-C): Serum was collected from WT (n=7) and MetS (n=8) mice and used to measure levels of (A) cholesterol, (B), glucose, and (C), triglycerides using an IDEX Catalyst One Analyzer. Compared to WT mice, MetS mice displayed significantly higher levels of (A) cholesterol (p=0.0003), (B) glucose (p=0.0003), and triglycerides (p=0.0011). Dotted lines indicate normal murine expression levels of these proteins. (D) To measure mRNA transcript expression of C reactive protein (Crp) in WT and MetS mice, blood RNA was isolated and qRT-PCR was utilized to plot expression normalized to GAPDH. MetS mice displayed significantly higher levels of Crp mRNA transcripts when compared to WT mice (p=0.0022). (E-L) To determine if MetS mice displayed signs of nonalcoholic fatty liver disease (NAFLD) progress to nonalcoholic steatohepatitis (NASH), liver lobes were collected from WT and MetS mice, fixed, embedded in paraffin, sectioned, and stained with hematoxylin and eosin (H&E) for pathological analysis by a blinded liver pathologist. Slides were then scored for the presence of (E) percent microsteatosis, (F) percent macrosteatosis, (G) total steatosis, and (H) percent hypertrophy. The incidence of each of these parameters was significantly higher in livers derived from MetS mice when compared to WT: microsteatosis (p=.0065), macrosteatosis (p=.0022), hypertrophy (p=.0022), and total steatosis (p=.0022). Panels (I-J) are representative liver H&E stains that highlight the criteria used for scoring in (E-H). Panel (I) shows normal liver histology of a WT mouse taken at 200x. Panel (J) highlights regions of macrosteatosis (solid triangle), and microsteatosis (dashed triangle) in a liver derived from a mouse with MetS. Panel (K) also highlights normal histology in a WT liver at 40x, compared to a region of cirrhosis (dashed and dotted triangle) noted in a MetS liver in Panel (L).

Further, we isolated RNA from blood collected from WT and high fat diet-fed mice to assess C reactive protein (CRP) levels relative to GAPDH via qRT-PCR. CRP is used as a serum biomarker to indicate acute inflammatory responses [36-38], and elevated baseline levels are linked to chronic inflammation due to tissue damage from excessive, unresolved inflammation (reviewed in [39]). CRP is an interesting biological marker to examine in our murine model as chronic inflammation is believed to be a triggering factor for MetS because various MetS-associated conditions, like obesity, enhance inflammatory cytokine secretion (reviewed in [40]).

Further, a human cohort study identified elevated CRP as an independent risk factor for mortality in SARS-CoV-2 patients hospitalized with COVID-19 [41]. As with the other biomarkers of MetS examined, **Figure 1D** identifies that high fat diet-fed mice displayed a significant elevation in baseline CRP transcripts when compared to WT counterparts (p=0.0022). In our previous publication, we noted that these high fat diet-fed mice displayed elevated levels of various pro-inflammatory cytokine-encoding mRNA transcripts [32], indicative of acute inflammation at baseline; coupling these data with the CRP levels presented here, high fed diet-mice show signs of chronic inflammation.

As we have previously reported elevated serum levels of normally liver-resident enzymes indicative of hepatocyte damage in our high fat diet murine model [32], we sought to investigate whether these animals showed pathological signs of non-alcoholic fatty liver disease (NAFLD). NAFLD describes the spectrum of liver disease where hepatic steatosis, or hepatocyte macrovesicular triglyceride accumulation, occurs independently from secondary causes like medication side effects or excessive alcohol consumption [42]. Important for our studies, NAFLD is also defined as the hepatic manifestation of MetS [43]. Liver lobes were collected from WT and high fat diet-fed mice after 12 weeks of respective diet feeding and were fixed, embedded in paraffin, sectioned, and stained with hematoxylin and eosin (H&E) for pathological analysis by a blinded liver pathologist. This pathologist adopted a murine NAFLD scoring model from a previous publication [44] to determine whether MetS mice displayed clinically relevant signs of liver pathology. Using this NAFLD activity score (NAS), percentages of hepatocytes displaying microsteatosis (**Figure 1E**) or macrosteatosis (**Figure 1F**) were recorded, and a total steatosis score (**Figure 1G**) was calculated utilizing these parameters. Further, livers were also scored for the presence of hypertrophic cells (**Figure 1H**). No signs of steatosis or hypertrophy were noted in WT mice, making the percentages of microsteatosis (p=.0065), macrosteatosis (p=.0022), and hypertrophy (p=.0022), in addition to the total steatosis score (p=.0022), significantly higher in high fat diet-fed mice when compared to WT counterparts (**Figure 1E-H**). **Figures 1I-J** show representative images of the criteria utilized for pathology scores from a WT (**Figure 1I**) and a high fat diet-fed (**Figure 1J**) mouse. When compared to the normal histology noted in the WT mouse liver depicted in **Figure 1I**, the dotted triangle in **Figure 1J** indicates a region of hypertrophy, while the solid triangle indicates macrosteatosis, and the dashed triangle indicates a region of microsteatosis in a high fat diet-fed mouse. Within the NAFLD spectrum of disease, nonalcoholic steatohepatitis (NASH) is an inflammatory subtype that can progress to cirrhosis [42]. While NASH affects 3-6% of the United States population, it is more prevalent in individuals with MetS, and progression to cirrhosis is associated with increased liver-specific as well as all-cause mortality [45]. In addition to showing signs indicative of NAFLD, several high fat diet-fed mice analyzed for pathology displayed signs of NASH which progressed to liver cirrhosis, as indicated by the solid-dashed triangle in **Figure 1L** when compared to the normal histology noted in **Figure 1K**. Taken together, these data suggest that after approximately 12 weeks of high fat diet feeding, mice display diagnostic criteria indicative of MetS. As such, this group of mice are referred to as MetS for the remainder of this study, with regular chow fed counterparts referred to as WT for wild type.

### Metabolic syndrome accelerates virally induced mortality and enhances inflammation at sites of viral replication

To interrogate the impact of MetS on SARS-CoV-2 infection outcomes, WT (n=5) and MetS (n=5) K18 mice were infected with 10^4^ FFU of an ancestral SARS-CoV-2 strain intranasally (IN). Mice were monitored and all MetS mice succumbed to infection by 9 DPI (**Figure 2A**). Although the WT K18s also uniformly succumbed to infection by 13 DPI, mortality was significantly accelerated in the MetS mice (p=0.0143) (**Figure 2A**). Cohort studies have revealed hyperinflammation as a contributing factor to worsened SARS-CoV-2 infection outcomes in COVID-19 patients with MetS (reviewed in [46]). Additionally, inflammatory cytokine storm has been linked to multi-organ failure in SARS-CoV-2 patients [47]. We therefore sought to determine whether MetS mice exhibited enhanced inflammation when compared to WT counterparts. To test this, WT (n=5) and MetS (n=5) K18 mice were infected with 10^4^ FFU of an ancestral SARS-CoV-2 strain IN. At 3 days post infection (DPI), mice were euthanized, organs were collected, and levels of the following inflammatory cytokines were measured in the spleen, lungs, and brain: tumor necrosis factor-α (TNF-α), interleukin-6 (IL-6), and interleukin-1β (IL-1β). In the spleen, mRNA transcripts were significantly higher in MetS mice for TNF-α (p=0.0159) and IL-1β (p=0.0317), but not for IL-6 (p=0.2222) (**Figure 2B**). In the lungs, mRNA transcripts were significantly higher in MetS mice for each cytokine tested: TNF-α (p=0.0079), IL-6 (p=0.0317), and IL-1β (p=0.0317) (**Figure 2C**). In brains derived from MetS mice, inflammatory cytokine levels were also significantly elevated for each cytokine tested: TNF-α (p=0.0317), IL-6 (p=0.0317), and IL-1β (p=0.0079) (**Figure 2D**). These data indicate that the MetS state fosters higher virally induced levels of inflammation at sites of viral replication, potentially contributing to the accelerated mortality noted in MetS mice when compared to WT mice.

**Figure 2:**
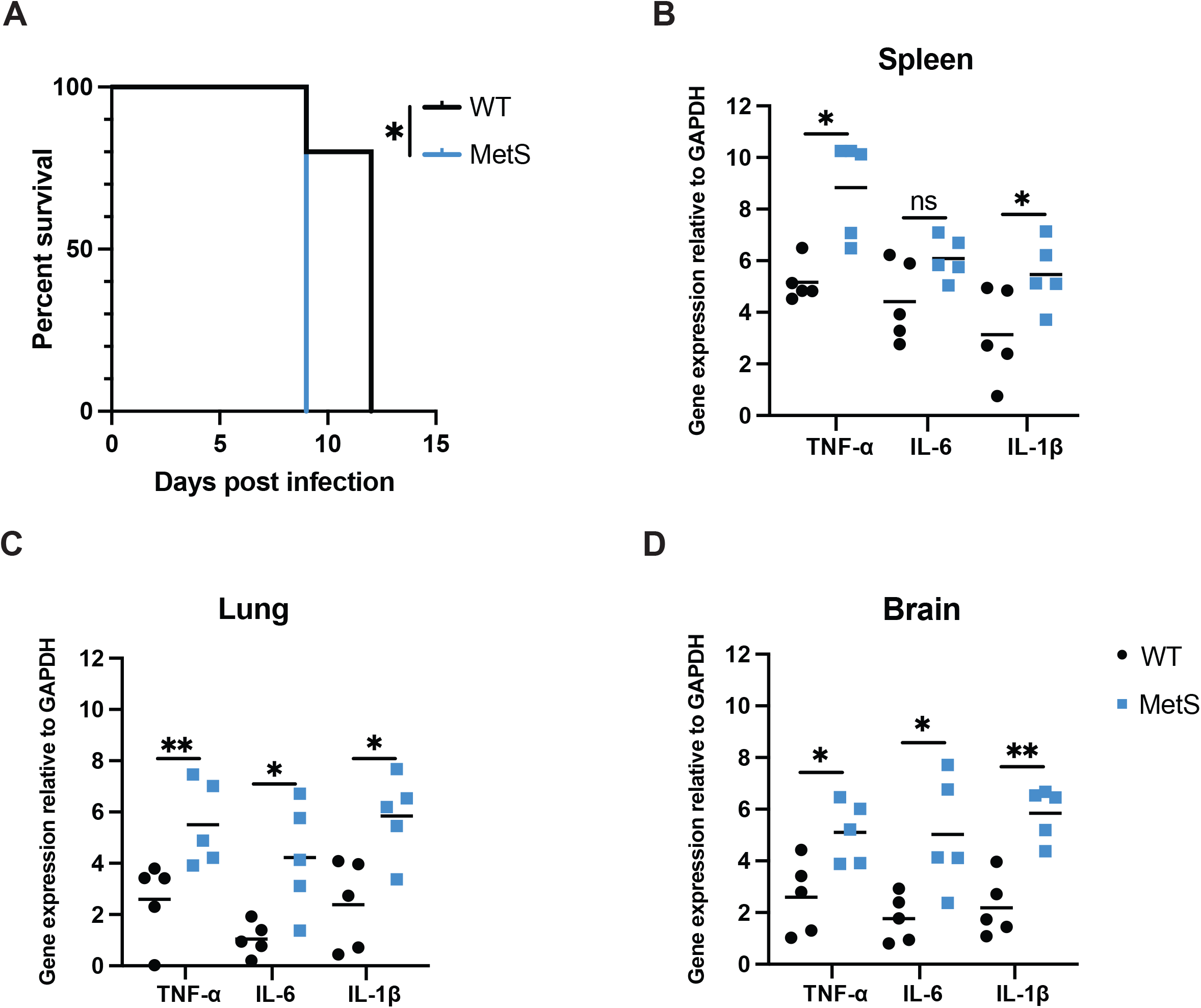
Metabolic syndrome accelerates virally induced mortality and enhances inflammation at sites of viral replication. (A) WT (n=5) and MetS (n=5) K18 mice were infected intranasally (IN) with 10^4^ focus forming units (FFU) of the ancestral SARS-CoV-2 strain. MetS mice showed significantly accelerated mortality (p=0.0143) when compared to survival of WT mice. (B-D) At 3 days post infection (3 DPI) expression of inflammatory cytokine mRNA was measured in the (B) spleen, (C) lung, and (D) brain of WT and MetS mice. To do this, RNA was isolated from organ homogenates and qRT-PCR was used to measure mRNA transcript levels relative to GAPDH expression in the same samples. In the spleen, MetS mice showed significantly higher levels of tumor necrosis factor-α (TNF-α) (p=0.0159) and interleukin-1β (IL-1β) (p=0.0317) transcripts, but no differences were noted between levels of interleukin-6 (IL-6) transcripts (p=0.2222). (C) In the lungs, TNF-α (p=0.0079), IL-6 (p=0.0317) IL-1β (p=0.0317) expression was significantly higher in MetS mice. (D) In the brain, expression levels of TNF-α (p=0.0317), IL-6 (p=0.0317), and IL-1β (p=0.0079) were significantly higher in MetS mice compared to WT counterparts.

### Metabolic syndrome influences viral replication patterns

After determining that the MetS state enhanced inflammation at sites of viral replication, we next sought to determine if viral titers differed between WT and MetS mice. To test this, WT (n=5) and MetS (n=5) K18 mice were infected with 10^4^ FFU of an ancestral SARS-CoV-2 strain IN. At 3 days post infection (DPI), mice were euthanized, organs were collected, and SARS-CoV-2 genome copies were measured in each organ using qRT-PCR. Viral titers were quantified utilizing a copy control. No significant differences in titer were noted in the spleen (p=0.8413) in terms of viral genome copies between WT and MetS mice (**Figure 3A**), but MetS mice displayed significantly higher levels of viral genome copies in the lungs (p=0.0079) (**Figure 3B**) and brain (p=0.0079) (**Figure 3C**) when compared to WT mice. Due to the significant elevation in SARS-CoV-2 genome copies in MetS lungs and brains, we also measured levels of infectious virus in these organs. No infectious virus was detected in the spleens of either WT or MetS mice at 3DPI (**Figure 3D**); however, there was a significantly lower amount of infectious virus found in the lungs (p=0.0079) (**Figure 3E**) and brains (p=0.0079) (**Figure 3F**) of MetS animals when compared to WT mice. These findings suggest that the MetS state fosters high levels of viral replication yet blunts the ability of infectious progeny from being assembled at primary sites of viral replication.

**Figure 3:**
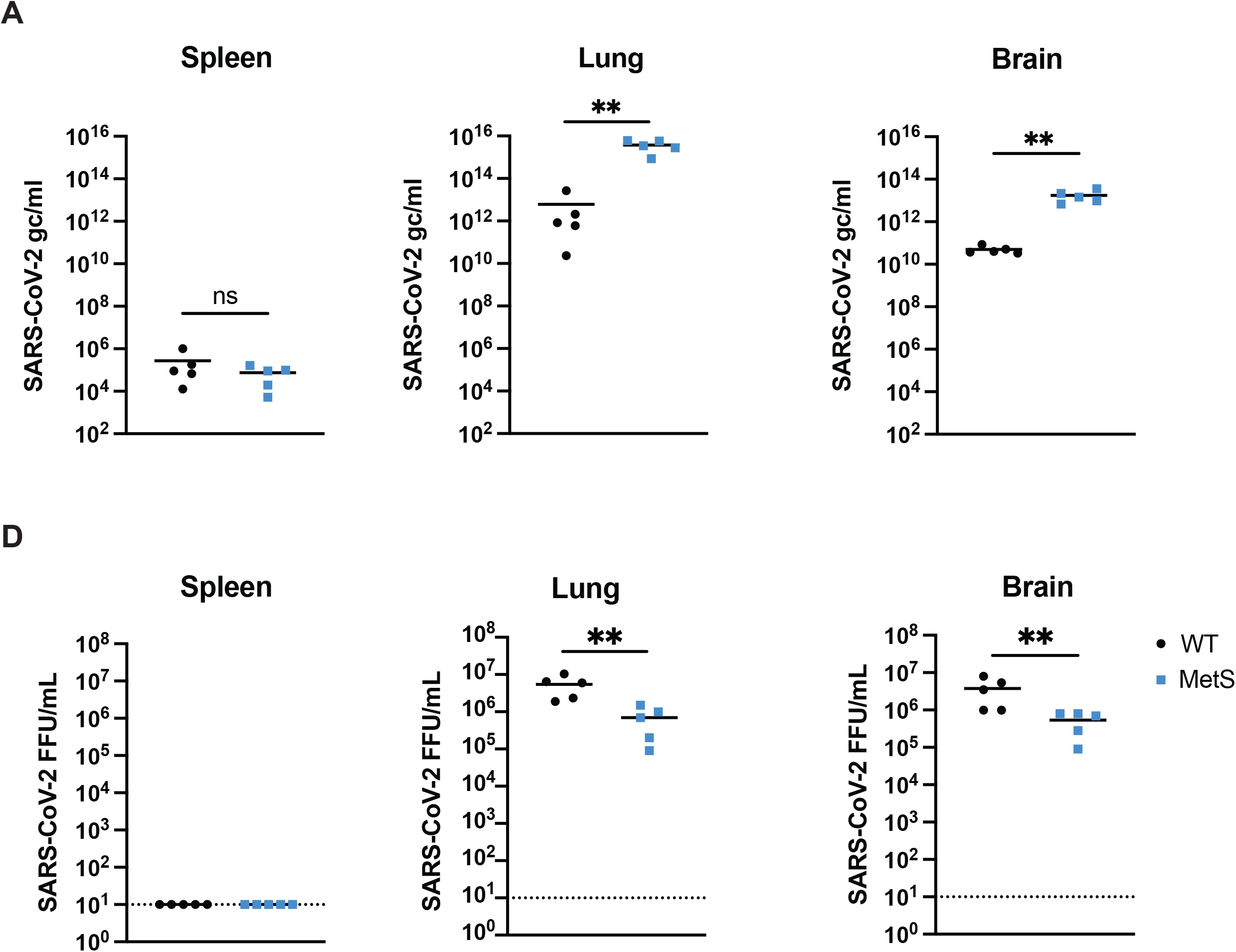
Metabolic syndrome influences viral replication patterns. (A-C) WT (n=5) and MetS (n=5) K18 mice were infected intranasally (IN) with 10^4^ focus forming units (FFU) of the ancestral SARS-CoV-2 strain. At 3 days post infection (DPI), RNA was isolated from the (A) spleen, (B) lung, and (C) brain of infected mice and qRT-PCR was utilized to measure viral genome copies quantified with a copy control. No significant differences were noted in viral genomes in the (A) spleen (p=0.8413), but genome copies were significantly higher in the (B) lung (p=0.0079) and (C) brain (p=0.0079) of MetS animals compared to WT. Focus forming assays were also performed on the (D) spleen, (E) lung, and (F) brain of WT and MetS mice. Infectious virus was not detected in the (D) spleen, but significantly lower amounts of infectious virus were detected in the (E) lung (p=0.0079) and (F) brain (p=0.0079) of MetS mice compared to WT mice.

### Metabolic syndrome impairs the efficacy of vaccine-conferred neutralizing antibodies

With reports of heightened SARS-CoV-2 disease severity in individuals with MetS [4-7], coupled with historical data linking obesity to impaired vaccine efficacy against hepatitis B virus, influenza viruses, SARS-CoV-2, and bacterial pathogens [48-60], we sought to interrogate the impact of MetS on vaccine-conferred neutralizing antibody responses in mice following immunization with the Moderna mRNA-1273 SARS-CoV-2 vaccine. As our data and previous reports have shown, K18 mice support high levels of SARS-CoV-2 replication in the brain [61-63], a site not reported to harbor much direct viral activity in humans [64], we instead utilized B6 mice where the course of SARS-CoV-2 infection is more similar to what is reported in humans. Additionally, with the evolution of viral variants, newer circulating SARS-CoV-2 strains can infect and induce illness in B6 mice [65, 66], eliminating the requirement for studies to be done exclusively in K18 mice.

To interrogate the impact of MetS on vaccine efficacy in B6 mice, we intramuscularly (IM) vaccinated WT (n=16) and MetS (n=14) mice with 5μg of the Moderna mRNA-1273 SARS-CoV-2 vaccine. At 8 days post vaccination (DPV), mice were submandibularly bled. Serum was collected from these vaccinated mice at various time points and used to analyze the neutralization capacity of antibodies primed in either WT or MetS mice. By utilizing focus reduction neutralization tests (FRNTs), we were able to determine the serum concentration, a proxy of neutralizing antibody titer, required to neutralize 90% (FRNT_90_) and 50% (FRNT_50_) of SARS-CoV-2 present in the assay.

We sought to determine if MetS impacted neutralizing antibody responses, as well as whether vaccine-conferred neutralizing antibody responses differed in their ability to neutralize the ancestral SARS-CoV-2 strain compared to a newer variant, Delta B.1.617.2. To test this, serum was collected from the mRNA-1273 vaccinated WT and MetS mice at 13 DPV. As can be seen in **Figure 4A**, there was an equivalent percentage of ancestral SARS-CoV-2 infected cells at each serum dilution tested for WT and MetS mice. Similarly, FRNT_90_ (p=0.1274) and FRNT_50_ (p=0.3543) values were equivalent between WT and MetS mice at 13 DPV against the ancestral SARS-CoV-2 strain (**Figures 4B** and **4C**, respectively). When examining the neutralization capacity of antibodies against the B.1.617.2 variant, both WT and MetS-primed neutralization antibodies exhibited a poorer capacity for neutralization compared to the ancestral strain of SARS-CoV-2 (**Figure 4D**). However, similar to neutralization against the ancestral strain, no differences were noted in the FRNT90 (p=0.4351) values (**Figure 4E**), but a reduction was noted in FRNT_50_ (p=0.0109) (**Figure 4F**) values of MetS-primed antibodies against B.1.617.2 when compared to WT-primed antibodies. As mentioned in the T cell studies, these mice were boosted at 21 DPV with a second mRNA-1273 dose. At 21 DPB, serum was collected from these animals and used for FRNTs again looking at neutralization of ancestral SARS-CoV-2 or the B.1.617.2 variant. Interestingly, MetS mice appear to have a defect in antibody longevity or recall as there is a higher percentage of SARS-CoV-2 infected cells at several of the serum dilutions tested when compared to WT mice (**Figure 4G**). Similarly, FRNT_90_ (p<0.0001) (**Figure 4H**) and FRNT_50_ (p<0.0001) (**Figure 4I**) values were significantly lower in MetS mice when compared to WT, highlighting an impairment in neutralization capacity. Further, similar to the data presented in **Figure 4D**, overall neutralizing antibody function is worse when neutralization of the B.1.617.2 variant (**Figure 4J**) is measured when compared to neutralization of ancestral SARS-CoV-2. Further, as can be seen when looking at the FRNT_90_ (p=0.0011) (**Figure 4K**) and FRNT_50_ values (p=0.0003) (**Figure 4L**), serum primed in MetS mice displayed a poorer neutralization capacity when compared to antibodies primed in WT mice.

**Figure 4:**
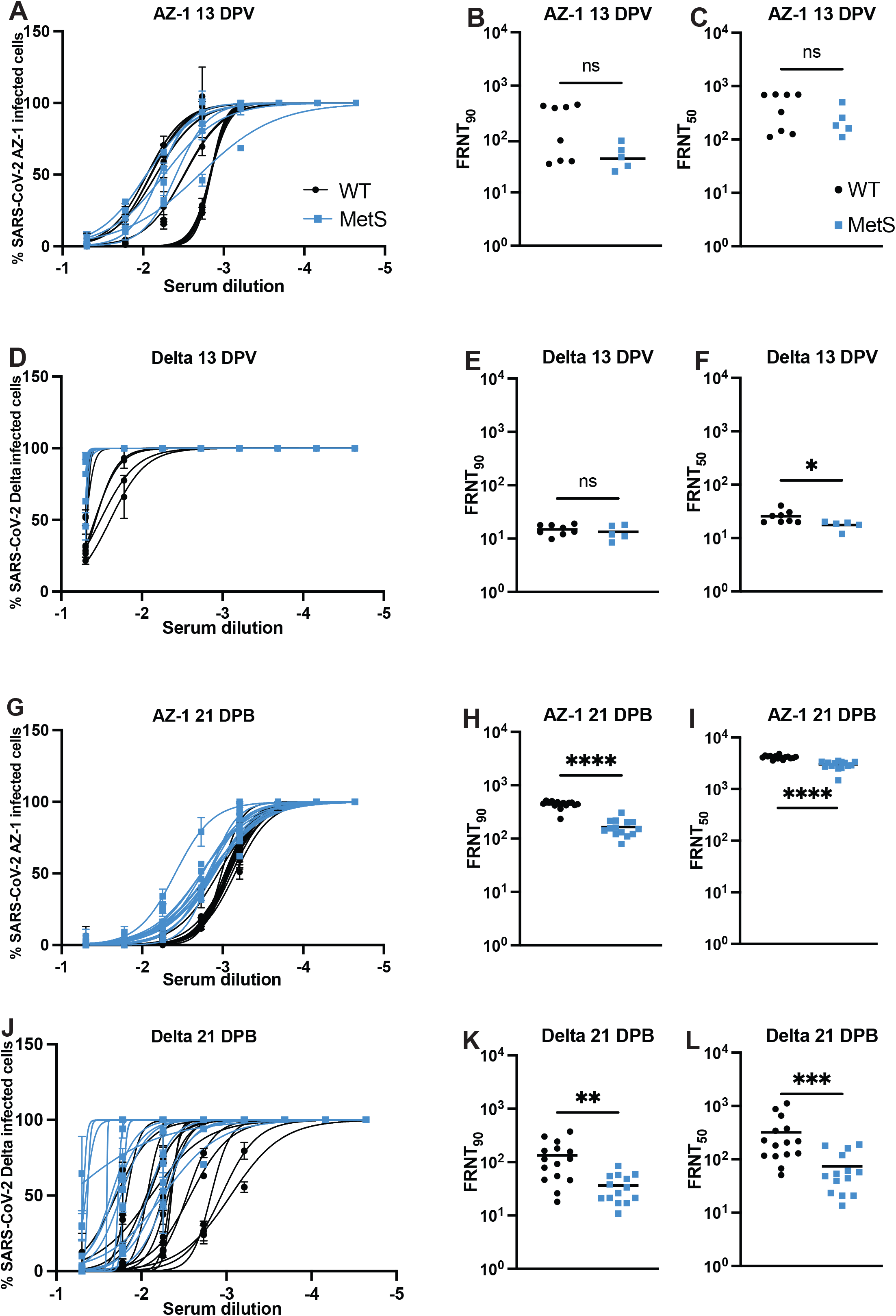
Metabolic syndrome impairs the efficacy of vaccine-conferred neutralizing antibodies. WT (n=15) and MetS (n=14) mice were vaccinated with the Moderna mRNA-1273 SARS-CoV-2 vaccine and boosted at 21 days post vaccination (DPV) with a second dose. Serum was collected from mice at (A-F) 13 DPV or 21 days post booster and used to perform focus reduction neutralization tests (FRNTs) to measure the ability of vaccine-conferred neutralizing antibodies to neutralize the (A-C, G-I) ancestral SARS-CoV-2 strain or the delta variant (D-F, J-L). At 13 DPV, the neutralization capacities of neutralizing antibodies primed in WT vs. MetS mice were virtually identical, as can be seen in the neutralization curve in panel A. (B) FRNT90 and (C) FRNT50 values were not significantly different (p=0.1274 and p=0.3543, respectively). Neutralization against the delta variant at (D) 13 DPV was worse in both groups of mice compared to their ability to neutralize the (A) ancestral strain, but no significant differences were noted in the (E) FRNT90 values between WT and MetS mice (p=0.4351). However, (F) FRNT50 values were significantly lower in MetS mice compared to WT counterparts (p=0.0109). At 21 days post boost, (G) neutralization was worse in MetS-primed antibodies compared to those primed in WT mice. (H) FRNT90 and (I) FRNT50 values were significantly lower in MetS mice compared to WT counterparts (p<0.0001 for both). As with 13 DPV, neutralization against the delta variant at (J) 21 days post boost was also worse in both groups of mice than against the (G) ancestral strain. Further, (K) FRNT90 and (L) FRNT50 values were significantly lower in MetS mice compared to WT (p=0.0011 and p=0.0003, respectively).

## Discussion

Because human cohort studies correlate MetS with impaired viral immunity and incidence of MetS is rising globally, we sought to characterize an animal model that would allow us to recapitulate this syndrome in mice to interrogate its impact on antigen-specific immunity. In the current study, we established and characterized a murine model of MetS which we utilized to study immune responses to SARS-CoV-2 infection and vaccination, as well as viral disease severity. We have previously illustrated that after approximately 12 weeks of high fat diet feeding, mice become obese, inflamed, and have liver dysfunction [32]. In the current study, we further characterized this model and concluded that high fat diet-fed mice also displayed hyperglycemia, hypertriglyceridemia, hypercholesterolemia, and elevated levels of CRP which indicate chronic inflammation (Figure 1A-D). In addition, these mice displayed liver pathology indicative of NAFLD progressed to NASH (Figure 1E-F). Taken together, these findings suggested that these mice developed MetS following prolonged high fat diet feeding.

Following viral infection, MetS K18 mice uniformly succumbed to SARS-CoV-2 at an accelerated rate when compared to WT counterparts (Figure 2A). The high mortality we noted while utilizing the K18 mouse model recapitulated the findings of others [61, 67, 68], but did limit our ability to study the adaptive arm of the immune system. The K18 model is superb for testing the efficacy of antivirals and vaccines against SARS-CoV-2 as they represent a stringent lethal challenge model. However, it is difficult to study other aspects of viral immunity following infection in these mice. For this reason, we recognize the benefit to utilizing B6 mice for characterizing aspects of viral immunity to SARS-CoV-2 infection, especially utilizing variants of concern that induce disease in B6 mice [65, 66]. Nonetheless, our findings suggest that the high inflammation displayed by MetS mice in the spleen, lung, and brain (Figure 2B-D) contributed to their accelerated mortality, as inflammatory cytokine storm has been widely reported in severe human cases of SARS-CoV-2 infection [46, 47, 69, 70].

Additionally, levels of viral genome copies were elevated in the same tissues in MetS mice that harbored high levels of inflammation (Figure 3A-C), potentially accelerating mortality in these mice. These data are intriguing as we anticipated that infectious titers would follow the same pattern as genome copy numbers, where MetS mice would have significantly elevated levels, but we found the opposite to be true (Figure 3D-F). Other murine studies utilizing high fat diet feeding models to explore the impact of obesity on viral immunity have noted discrepancies in terms of viral titer differences, with some studies showing viral load to be higher in mice fed a high fat diet compared to those fed a regular chow diet [54, 55, 71], while other studies reported lower titers in high fat diet fed mice [72, 73]. Our findings suggest that the time post infection is critical to understanding the discrepancies regarding viral load differences in high fat diet-fed mice. It is possible that titer differences are often noted at early time points post infection due to a blunted and delayed type I IFN response in mice fed a high fat diet [54, 55, 74, 75], thus resulting in an elevated viral load. It is therefore conceivable that this blunted IFN response could contribute to the impaired adaptive immunity associated with metabolic dysfunction. Interestingly, the ability for SARS-CoV-2 to induce a pathological effect in hosts has been linked to the virus’s ability to antagonize type I IFN responses [76-79]. As such, it is possible that the propensity for metabolic perturbances to blunt type I IFN responses coupled with virally-induced delayed type I IFN responses compound in the MetS mice, thus elevating early viral disease severity.

Following vaccination, MetS mice displayed similar neutralizing antibody responses against the ancestral SARS-CoV-2 strain when compared to WT mice (Figure 4A-C) and although overall neutralization against the delta variant was worse in both groups of mice when compared to neutralization of the ancestral strain (Figure 4D-F), MetS mice displayed significantly worse FRNT_50_ values against the delta variant following primary vaccination. Further, MetS mice mounted significantly worse neutralizing antibody responses post-booster dose against both the ancestral SARS-CoV-2 strain (Figure 4G-I) and the delta variant (Figure 4 J-L) when compared to WT mice. These findings are consistent with human studies following influenza vaccination where individuals with obesity mounted an initial robust antibody response, yet antibody titers in individuals with obesity waned more rapidly than in normal weight individuals [58]. Our findings, coupled with previous work, support a role for MetS-associated comorbidities in impairing the ability of vaccine-conferred immune responses to persist in a memory population that can be recalled during a future antigenic encounter. Further, the overall reduction in the ability of MetS-primed antibodies to neutralize SARS-CoV-2 infection supports additional findings of human cohort analyses. For example, one study found a significant reduction in SARS-CoV-2 antibody levels in individuals with obesity following infection compared to individuals of a healthy weight, and the authors noted that a majority of these antigen-specific antibodies were non-neutralizing and auto-reactive [80]. Other studies noted titers of SARS-CoV-2-specific antibodies to be lower in individuals with obesity compared to those without [81]. We found it interesting that even following primary vaccination at 13 DPV, MetS mice already showed an impaired ability to neutralize the delta variant when compared to WT mice but did not show an impairment in their ability to neutralizing the ancestral SARS-CoV-2 strain. This finding likely reflects the ability of the delta variant to escape a degree of vaccine-conferred immunity [82-85], but calls closer attention to the potential for enhanced viral escape of the delta variant in individuals with MetS.

These studies enabled us to conclude that mice fed a high fat diet for at least 12 weeks experienced the same MetS-associated comorbidities as human MetS patients. Our infection studies revealed that MetS accelerates mortality to SARS-CoV-2 and enhances expression of inflammatory cytokine mRNA at sites of infection. Further, MetS promoted a high amount of viral replication in the lungs and brain yet seemed to inhibit the formation of infectious viral progeny at these tissue sites. Finally, MetS blunted the neutralizing antibody response induced by vaccination, particularly following the booster vaccine. Future studies to identify the mechanism(s) by which the MetS state blunts the neutralization capacity of neutralizing antibodies are critical for furthering this field and identifying ways to improve viral immunity among individuals with MetS.

## Materials and Methods

### Ethics Statement

All animal studies were conducted in accordance with the National Institutes of Health Guide for Care and Use of Laboratory Animals and approved by both the University of Kentucky and the Saint Louis University Animal Care and Use Committees.

### Virus and Cells

SARS-CoV-2 ancestral strain (Isolate USA-AZ1/2020 BEI NR-52383) and Delta strain were each passaged once in Vero cells (African green monkey kidney epithelial cells) transfected to express human ACE2 and TMPRSS2, hereafter referred to as VAT cells. Viruses were titered *via* focus forming assays on VAT cells as previously described [86].

### Mice

C57BL/6J (B6) and B6.Cg-Tg(K18-ACE2)2Prlmn/J (K18) were purchased commercially from Jackson Laboratories and housed in a pathogen-free mouse facility at the University of Kentucky or Saint Louis University School of Medicine. 3-week-old mice were fed either a control (WT) or high fat diet (40% kcal fat, 20% kcal fructose and 2% cholesterol, Research Diets Inc.) for approximately 12 weeks. At 12 weeks of respective diet feeding, high fat diet-fed mice were analyzed for signs of MetS and utilized in our studies once deemed to have MetS.

### Measurement of MetS parameters

Serum was collected from WT (n=7) and MetS (n=8) B6 mice that had been fed their respective diets for 12 weeks. Serum was diluted 1:2 in 1X PBS and loaded into sample collection cups for analyses utilizing an IDEXX Catalyst One Chemistry Analyzer. Clips were loaded into the machine to measure serum levels of cholesterol, glucose, and triglycerides. To measure levels of C reactive protein, blood was collected from WT (n=6) and MetS B6 (n=6) mice fed their respective diets for 12 weeks. Blood was collected into tubes containing RNAzolBD (Molecular Research Center, Inc.: RB 192), and RNA was isolated according to the manufacturer’s protocol. mRNA expression of C reactive protein was measured through qRT-PCR using a Taqman primer probe set purchased from Integrated DNA Technologies (IDT) based on the following assay identifier: Mm.PT.58.46041922. Relative expression was determined by 2^ΔΔCT^ with fold induction relative to GAPDH (assay identifier: Mm.PT.39a.1) expression for the same sample. To examine liver pathology indicative of nonalcoholic fatty liver disease (NAFLD), liver lobes were collected from WT (n=6) and MetS (n=6) B6 mice following perfusion with paraformaldehyde (PFA). Lobes were collected into a solution of 4% PFA, then transferred to 1X PBS 24 hours later. Lobes were then paraffin embedded, sectioned, mounted onto slides, and stained using hematoxylin and eosin (H&E) to visualize tissue morphology. Slides were given to a licensed liver pathologist and scored blindly based on the following criteria: percent of tissue displaying hypertrophic cells, microsteatosis, or macrosteatosis, and samples were given a total steatosis score, as well. Tissues were also examined for signs of cirrhosis indicative of NAFLD progressed to nonalcoholic steatohepatitis (NASH).

### Viral infections

For studies examining acute viral infection outcomes, WT (n=5) and MetS (n=5) K18 mice were infected with 10^4^ FFU of SARS-CoV-2 AZ-1 intranasally (IN). For studies examining T cell responses to viral infection, WT (n=5) and MetS (n=5) K18 mice were infected with 10^4^ FFU of SARS-CoV-2 AZ-1 IN.

### Measurement of inflammatory cytokine mRNA expression

15-week-old WT (n=5) and MetS (n=5) K18 mice were infected IN with 10^4^ FFU SARS-CoV-2 AZ-1. At 3 DPI, intracardiac perfusion with 20ml of 1X PBS was performed and spleens, lungs, and brains were collected. These tissues were homogenized in DMEM supplemented with 5% FBS using a BeadMill 24 (Fisher Scientific). RNA was extracted from organ homogenates using TriReagent RT (Molecular Research Center Inc.: RT111). mRNA expression of tumor necrosis factor-α (TNF-α), interleukin-1β (IL-1β), and interleukin-6 (IL-6) was determined using qRT-PCR with Taqman primer probe sets purchased from Inegrated DNA Technologies (IDT) based on the following assay identifiers: Mm.PT.58.12575861 (TNF-α), Mm.PT.58.41616450 (IL-1β), and Mm.PT.58.10005566 (IL-6). Relative expression of each cytokine was determined by 2^ΔΔCT^ with fold induction relative to GAPDH (assay identifier: Mm.PT.39a.1) expression for the same sample.

### Measurement of viral titers

15-week-old WT (n=5) and MetS (n=5) K18 mice were infected IN with 10^4^ FFU SARS-CoV-2 AZ-1. At 3 DPI, intracardiac perfusion with 20ml of 1X PBS was performed and spleens, lungs, and brains were collected. Organs were homogenized in DMEM supplemented with 5% FBS using a BeadMill 24 (Fisher Scientific). RNA was extracted from organ homogenates using TriReagent RT (Molecular Research Center Inc.: RT111). SARS-CoV-2 genome copies were quantified via qRT-PCR using a primer probe set purchased from IDT with the following sequence: Forward 5’ GAC CCC AAA ATC AGC GAA AT 3’, Reverse 5’ TCT GGT TAC TGC CAG TTG AAT CTG 3’, Probe 5’ ACC CCG CAT TAC GTT TGG TGG ACC 3’. Viral genome copies were quantified based on a standard curve generated from a SARS-CoV-2 copy control (purchased from BEI).

### Vaccination

15-week-old male and female WT (n=15) and MetS (n=14) B6 mice were intramuscularly (IM) vaccinated with 5μg of the Moderna mRNA-1273 vaccine. At 21 days post vaccination (DPV), mice were IM boosted with a second 5μg dose of the Moderna mRNA-1273 vaccine.

### Focus reduction neutralization tests (FRNTs)

15-week-old male and female WT (n=15) and MetS (n=14) B6 mice were intramuscularly (IM) vaccinated with 5μg of the Moderna mRNA-1273 vaccine. At 13 DPV, serum was collected from vaccinated mice and stored. At 21 days post booster vaccine, serum was again collected and stored. Using serum from the aforementioned time points, FRNTs were conducted to measure neutralizing antibody function as previously described [87]. Briefly, serum was diluted in DMEM supplemented with 5% FBS and combined with ∼100 FFU of either SARS-CoV-2 AZ-1 or Delta. Serum:virus samples were allowed 1 hour at 37°C, 5% CO_2_ to form immune complexes in 96-well round bottom plates. Antibody:virus complexes were then transferred to 96-well flat bottom plates pre-seeded with a monolayer of VAT cells. After a 1 hour incubation at 37°C, 5% CO_2_, cells were overlaid with 2% methylcellulose warmed to room temperature and diluted in DMEM supplemented with 5% FBS. Plates were returned to the incubator and allowed to rest for 24 hours. Following infection, cells were fixed with 5% PFM in 1X PBS for 15 minutes at room temperature. Plates were further decontaminated in 5% PFM and washed in 1X PBS prior to being removed from BSL-3 containment. Cells were then washed with 1X PBS 3 times and permeabilized at room temperature for 10 minutes with 0.05% Triton-X in 1X PBS. Infected VAT cell foci were stained with anti-SARS polyclonal guinea pig sera (BEI) 1:1000 overnight at 4°C. Cells were then washed 3 times with 0.05% Triton-X in 1X PBS and stained 1:5000 with horseradish peroxidase-conjugated goat anti-guinea pig IgG for 2 hours at room temperature. Cells were washed again 3 times with 0.05% Triton-X in 1X PBS, and TrueBlue KPL peroxidase substrate was added to each well to visualize infected foci as blue spots. Foci were counted using an ImmunoSpot CTL Elispot plate reader.

## Author Contributions

EG and AKP conceptualized the work, wrote, and edited the manuscript. EG, AKP, and JDB aided in experimental design. DC conducted liver histopathological analysis. EG, JDB, ETS and AMD completed the FRNTs. ETS established and maintained K18 mouse colony. Execution of all other experiments was performed by EG. AKP was responsible for the support of experiments. All authors contributed to the article and approved the submitted version.

## Acknowledgments

We would like to thank Dr. Mariah Hassert for support in protocol generation and developing the SARS-CoV-2 infection model in the laboratory.

